# *In vitro* and *in vivo* activity profiles of broad-spectrum bacterial ATP synthase inhibitors

**DOI:** 10.1101/2025.08.21.671483

**Authors:** Suzanne Chamberland, Audrey Larose, Guillaume Millette, Jean-Philippe Langlois, Julie Côté-Gravel, Eric Brouillette, Daniela Droppa Almeida, Julien A Delbrouck, Iryna Diachenko, Abdelkhalek Ben Jamaa, Chad Normandin, Alexandre Murza, François Malouin, Pierre-Luc Boudreault

**Affiliations:** Département de biologie, Faculté des sciences, Université de Sherbrooke, Sherbrooke, J1K 2R1, Québec, Canada; Département de Pharmacologie-Physiologie, Faculté de Médecine et des Sciences de la Santé, Université de Sherbrooke, Sherbrooke, J1H 5N4, Québec, Canada; Institut de Pharmacologie de Sherbrooke, Sherbrooke, J1H 5N4, Québec, Canada

**Author notes:** **Corresponding authors:** Pierre-Luc Boudreault (Chemistry), François Malouin (Microbiology).

**Keywords:** ATP synthase inhibitors, broad-spectrum antibiotics, steroidal alkaloid, Gram-negative

## Abstract

Tomatidine (**TO**), a natural steroidal alkaloid derived from Solanaceae (e.g., tomato), is recognized for its narrow-spectrum antibiotic activity, particularly against persistent forms of *Staphylococcus aureus*, such as small-colony variants (SCVs). Previous studies have shown that **TO** exerts its effect by inhibiting *S. aureus* ATP synthase. In earlier work, we synthesized nearly 100 **TO** analogs featuring an ethylenediamine linker branched with aromatic substituents at the C3 position, and demonstrated that several of these analogs possess notable antibacterial activity against typical (non-SCV) *S. aureus* strains, including methicillin-resistant *S. aureus* (MRSA), with minimum inhibitory concentrations (MICs) ranging from 1 to 4 µg/mL. Among these, analogs incorporating an indole (**TM-247**) or para-substituted aryl moiety (**TM-184**, -I; **TM-218**, -Cl; **TM-220**, -Br; **TM-303**, -CF₃) emerged as lead candidates, exhibiting potent antibacterial activity against both Gram-positive and Gram-negative bacteria, including *S. aureus* SCVs. In the present study, we conducted a comprehensive antibacterial profiling of this compound series, including the predecessor compound **TM-02**, against a panel comprising 16 Gram-positive strains, 16 antibiotic-resistant *Escherichia coli* isolates, and two multidrug-resistant *Acinetobacter baumannii* strains. **TM-184** emerged as the most promising candidate and was subsequently subjected to an expanded *in vitro* evaluation across 24 clinically relevant Gram-negative bacterial species. Furthermore, **TM-184** was assessed *in vivo* using a neutropenic mouse thigh infection model against *E. coli*, where it demonstrated significant efficacy, leading to a substantial reduction in bacterial burden.

## 1. Introduction

Antibiotic resistance in bacterial pathogens has been a longstanding concern within the scientific community. However, it wasn’t until the O’Neill report [1] projected alarming mortality rates for 2050 that the general public and government authorities began to recognize the severity of this threat to human health. To enhance preparedness, the World Health Organization [2] 2017 identified a list of critical bacterial pathogens urgently needing new treatments. This list included Methicillin-resistant *Staphylococcus aureus*, as well as carbapenem-resistant Gram-negative *Enterobacteriaceae* and *Acinetobacter baumannii*. Notably, the latter two pathogens produce a β-lactamase that hydrolyzes carbapenems, often the last line of defense against these infections [3]. Additionally, the horizontal transfer of antibiotic resistance genes among bacteria has dramatically diminished the effectiveness of all current antibiotic classes. Newly developed antibiotics from these classes quickly become ineffective due to these resistance mechanisms [4,5]. Currently, only a few genuinely novel antibiotic molecular scaffolds target new mechanisms in the drug and clinical development pipelines [6,7]. We propose C3-derivatives of the steroidal alkaloid tomatidine (**TO**) as a novel antibiotic molecular scaffold. **TO**, a natural product with well-documented biological activities [8], exhibits potent narrow-spectrum antibiotic activity, particularly against respiratory-deficient small colony variants (SCVs) of *Staphylococcus aureus*. Our previous research demonstrated that SCVs’ heightened susceptibility to **TO** is due to its ability to inhibit bacterial ATP synthase, leading to the production of reactive oxygen species [9,10]. Bacterial ATP synthases are underutilized molecular targets, highlighted by the therapeutic relevance of bedaquiline, the first clinical ATP synthase inhibitor approved for treating *Mycobacterium tuberculosis* [11,12]. Despite bedaquiline’s cardiotoxicity linked to off-target effects [13], there remains potential for developing new ATP synthase inhibitors with different molecular scaffolds.

In an earlier study, we demonstrated that modifying the C3-position of tomatidine (**TO**) could yield ATP synthase inhibitors effective against *E. coli*. Specifically, derivatives with an ethylene diamine appendage and an aromatic substituent at the C3 position of the steroidal backbone showed enhanced outer membrane permeability and greater affinity for the *E. coli* ATP synthase, as confirmed by *in silico* docking experiments [14]. This study provides an in-depth investigation of the antibacterial activity profiles of the most promising C3-derivatives of tomatidine (TO), selected from a library of 100 analogs previously evaluated, including TM-247 bearing an indole, and para-substituted analogs (TM-184, I; TM-220, Br; TM-303, CF₃) [14]. These compounds exhibit broad-spectrum antibacterial activity against several of the most challenging pathogens, including carbapenem-resistant *E. coli*, extreme drug-resistant *A. baumannii*, methicillin-resistant *S. aureus* (MRSA) and vancomycin-resistant *Enterococcus faecium* (VRE). Through comprehensive *in vitro* and *in vivo* studies, this investigation offers a detailed examination of the antibacterial spectrum of these new bacterial ATP synthase inhibitors, thus highlighting the potential of these molecules as effective treatments in the ongoing battle against antibiotic-resistant infections.

## 2. Results

### 2.1. Synthesis of tomatidine analogs

We have previously reported the synthesis of the C3-modified **TO**-analogs assessed in this manuscript [14]. Briefly, starting from **1,** *N*,*O*-double formylation was carried out using acetic formic anhydride and Hünig’s base in tetrahydrofuran. This was followed by a chemoselective hydrolysis of the carbonate group, resulting in the *N*-formyltomatidine intermediate **2**. Oxidation with Dess-Martin periodinane provided 3-keto-*N*-formyl-tomatidine **3** with 88% yield (**Scheme 1**, **Panel A**). A subsequent reductive amination with ethylenediamine led to the final compound FC04-100 (**TM-02**). The reductive amination was non-stereoselective and proceeded under acid-catalysis, producing an equimolar mixture of diastereoisomers. Substituted bis-amino alkyl derivatives BocNH(CH-**R**)CH_2_NH_2_ were synthesized from their respective D-Boc protected amino acids following the synthetic route depicted in the dotted frame. First, *N*-Boc residues **4** were converted into primary amides **5** by adding ethyl chloroformate and triethylamine in tetrahydrofuran. Borane dimethyl sulfide complex in tetrahydrofuran was used to reduce amides, affording the respective amino-alkyl carbamate **6** (**Scheme 1**, **Panel B**). These intermediates were then subjected to reductive amination without any further purification, resulting in indole (**TM-247**), and para-aryl derivatives (X = -I, **TM-184**; -Br, **TM-218**; -Cl, **TM-220**; -CF_3_, **TM-303**). Finally, the *N*-formyl and *N*-Boc protecting groups were simultaneously removed under acidic conditions to provide C-3 **TO**-analogs.

**Scheme 1.**
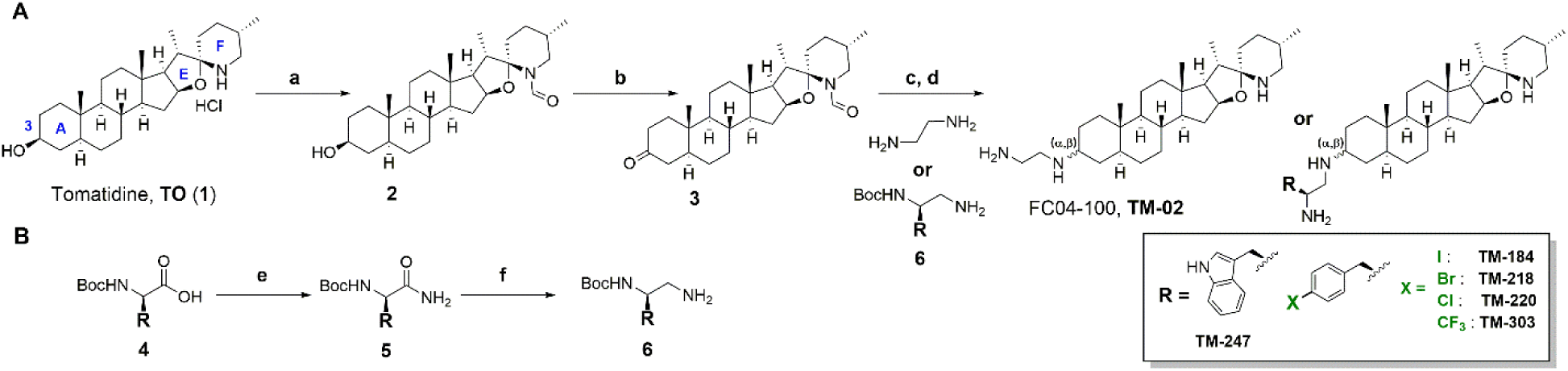
Structure and synthetic route of TM-compounds. *^a^*Reagents and conditions: Panel A: (a) i) acetic formic anhydride, DIPEA, THF, rt; ii) Na_2_CO_3_ buffer pH = 9.5, rt; (b) Dess-Martin periodinane, DCM, rt, 88% over two steps; (c) H_2_N-(CH_2_)_2_-NH_2_ or BocNH(CH-**R**)CH_2_NH_2_, AcOH_cat_, NaBH(OAc)_3_, MeOH, pH = 6, reflux; (d) AcCl, MeOH, 65°C, 5-40% over two steps. Panel B: (e) ClCO_2_Et, Et_3_N, THF, 0°C then NH_4_Cl_aq_, 0°C (quant.); (f) BH_3_.Me_2_S, THF, rt. 30-80%.

### 2.2. Specific *in vitro* biological activities of TO analogs

In a research program aimed at evaluating the potential of steroidal alkaloids as novel antibacterial agents, we previously synthesized and biologically assessed compounds in which structural modifications focused on the spiroaminoketal moiety (E, F rings) as well as on the screening of C3-appendages [14, 15]. **TM-02** was the first analog showing significant activity against prototypical, non-SCV, *S. aureus* (MIC of 4-8 µg/ml depending on the strain) but also having a moderate potency against *E. coli* (MIC of 32 µg/ml). In this study, we thoroughly investigated the antibacterial potential of **TM-02** along with five other **TM**-analogs (**Scheme 1**), which were discovered recently through a structure-activity relationship (SAR) study [14].

**Table 1** shows that the MICs of the C3-derivatives of **TO** changed from ≤0.25 (ATCC 29740*hemB*) to 1-2 µg/ml against a *S. aureus* SCV having a point mutation in *atpE* (ATCC 29740*hemB-atpE* G18C), as previously reported for the first derivative **TM-02** (*aka* FcM or FC04-100) [9]. The SCV *atpE*G18C was selected by serial passage on **TO** [9] and showed complete resistance to **TO** (MIC >128 µg/ml). This significant change in MIC establishes a clear link between the whole-cell activity of C3-derivatives and the bacterial ATP synthase target, although the loss of activity for the *atpE*G18C mutant was not as substantial as that observed for **TO** (**Table 1**). Besides, the C3-derivatives of **TO** now having Gram-negative antibacterial activity, prompted an investigation of the relative contribution of efflux and permeability in the overall activity of the selected **TM**-compounds against *E. coli*. For such a purpose, we used an AcrAB efflux pump mutant, which is hypersusceptible to compounds that are normally effluxed, as well as strain *E. coli lptD4213*, which has a compromised outer membrane barrier, and thus, is hypersusceptible to compounds that are unable to cross the outer membrane. **Table 1** presents the MIC ratios of the parental strain (strain MC4100 WT) over those of the mutants. Compounds **TM-02** and **TM-184** were not significantly affected by efflux, with a ratio of 2, while **TM-247** and **TM-303** were more affected (ratio of 4). In contrast, permeability ratios were higher for compounds with lower activity against *E. coli* (*i.e.*, higher MIC), namely **TM-02**, **-247**, and **-303**, with 32, 8, and 8 ratios, respectively. **TM-184** was the least affected by the permeability barrier, with a ratio of 4. Notably, despite an apparent reduced affinity for the *E. coli* ATP synthase compared to **TM-02**, as also observed for the *S. aureus* enzyme (**Table 1**, IC_50_), **TM-247** and **TM-303** exhibited stronger or equivalent activity against *E. coli* WT strains. This is likely attributed to their enhanced ability to cross the outer membrane barrier compared to compound **TM-02**, as concluded from the permeability ratios, given their higher hydrophobicity.

**Table 1.** *In vitro* biological activities of TM-analogs against *S. aureus* and *E. coli*.

|  | Broth microdilution MIC (µg/ml) |  |  |  |  |
| --- | --- | --- | --- | --- | --- |
|  | TO | TM-02 | TM-184 | TM-247 | TM-303 |
| **Bacterial strain** |  |  |  |  |  |
| *S. aureus* |  |  |  |  |  |
| ATCC 29740 (Newbould) | >128 | 4 | 2 | 2 | 4 |
| ATCC 29740*hemB* | ≤0.25 | ≤0.25 | ≤0.25 | ≤0.25 | ≤0.25 |
| ATCC 29740*hemB-atpE* G18C | >128 | 2 | 1 | 2 | 2 |
| *E. coli* |  |  |  |  |  |
| ATCC 25922 | >128 | 32 | 4 | 8 | 8 |
| MC4100 WT | >128 | 16 | 8 | 16 | 16 |
| MC4100 *acrAB* | >128 | 8 | 4 | 4 | 4 |
| MC4100 *lptD4213* | >128 | 0.5 | 2 | 2 | 2 |
| **MIC ratio** |  |  |  |  |  |
| **Efflux** (MIC MC4100 WT/*acrAB*) |  | 2 | 2 | 4 | 4 |
| **Permeability** (MIC MC4100/*lptD4213*) |  | 32 | 4 | 8 | 8 |
| **IC<sub>50</sub> (µM)** |  |  |  |  |  |
| **Bacterial ATP synthase** |  |  |  |  |  |
| *S. aureus* ATCC 29213 |  | 0.51 ± 0.17 | 0.70 ± 0.33 | 1.06 ± 0.24 | 1.32 ± 0.22 |
| *E. coli* ATCC 25922 |  | 1.49 ± 0.28 | 1.57 ± 0.17 | 2.12 ± 0.32 | 3.15 ± 0.73 |
| **Selectivity index** |  |  |  |  |  |
| **Hep2 Mitochondria ATP synthase** |  |  |  |  |  |
| Ratio IC<sub>50</sub> TM-compound/Oligomycin |  | 1652 | 845 | 1366 | 978 |

It is important to note that the acknowledged two-fold variability inherent in MIC assays complicates the interpretation of such data. However, the MIC values presented in **Table 1** underwent multiple confirmations. Overall, this analysis indicates that enhancing permeability could enhance the activity (MIC) of **TM-02** derivatives against *E. coli* by a factor of 4 to 8. There is also a possibility, to some extent, that a reduction in efflux may have contributed to this improvement. However, this was achieved through structural modifications that apparently slightly impaired target affinity in the case of **TM-247** and **TM-303** (slightly higher IC_50_ values, **Table 1**). **TM-184** was thus identified as the most promising compound. Using enzyme kinetics assays, we demonstrated that this compound could inhibit the *E. coli* ATP synthase activity with a K_i_ of 0.8-1.3 µM (0.55-0.89 µg/ml), which was comparable to that previously determined for **TM-02** (1.1 µM or 0.51 µg/ml) [16], or to the K_i_ of our positive control 4-chloro-7-nitrobenzofurazan (NBD-Cl). NBD-Cl is known to be a potent ATP synthase inhibitor [17], showing a Ki 1.8 µM in our assay. IC_50_ graphs for **TM-184** using both *S. aureus* and *E. coli* membrane vesicles are shown in **Figure S1**, and enzyme kinetics inhibition for TM-184 and NBD-Cl are shown in **Figure S2** (see supplementary materials).

Finally, we assessed the impact of structural modifications on the relative selectivity of **TM**-analogs. To gauge this, we determined the IC_50_ values of the compounds for mammalian mitochondria isolated from Hep2 cells, comparing them to oligomycin, a widely employed control drug known for inhibiting the mammalian ATP synthase enzyme [18]. In our assay, oligomycin demonstrated an IC_50_ of 0.014 µM. Consequently, we calculated a selectivity index, defined as the ratio of the IC_50_ of a specific **TM**-compound to that of oligomycin for Hep2 mitochondria. The results, presented in **Table 1** (selectivity index: ratio IC_50_ **TM**-compound/oligomycin), revealed that the **TM**-analogs were at least 800-fold less potent than oligomycin against the eukaryotic enzyme. Notably, **TM-184** exhibited the lowest selectivity. Whether this is attributable to its affinity for the enzyme or its ability to penetrate cells remains a subject for further investigation. Supplementary material **Figure S3** provides a comparison of ATP production by mitochondria in the presence of **TM-184** and oligomycin.

### 2.3. Spectrum of activity

The spectrum of activity of **TO** analogs, and more specifically of **TM-184**, was evaluated in greater depth. **TM-184** was also compared to other closely related derivatives to evaluate the effect of various halogen substitutions on the activity (**Scheme 1**, **TM-184**, Iodine; **TM-218**, Bromine; and **TM-220**, Chorine). The series demonstrated consistent antibacterial activity against Gram-positive bacteria, with MICs ranging from 1 to 4 µg/ml for *Staphylococcus spp*. (coagulase positive and negative) and *Enterococcus faecium* (**Table 2**). The strains tested included methicillin-resistant staphylococci (MRS) as well as vancomycin-resistant *Enterococcus* (VRE). *Bacillus spp*., *Listeria monocytogenes* and groups A and B streptococci appeared slightly less susceptible to some of the **TO** derivatives however **TM-184** consistently showed MIC ranging from 2 to 4 µg/ml. The nature of the halogen did not consistently impact the antibacterial activity. Hence, this analysis showed that the structural variations that modulated whole-cell activity against *E. coli* and the slight differences in the relative affinity to the ATP synthase enzyme (**Table 1**) had little effect on the overall activity against the Gram-positive microorganisms lacking an outer membrane barrier.

**Table 2.** *In vitro* activity profile of TM-analogs against Gram-positive bacteria.

| Bacterial strain | Broth microdilution MIC (µg/ml) |  |  |  |  |  | CIP | MEM | VAN |
| --- | --- | --- | --- | --- | --- | --- | --- | --- | --- |
|  | TM-02 | TM-184 | TM-218 | TM-220 | TM-247 | TM-303 |  |  |  |
| *Staphylococcus aureus* ATCC 29213 | 8 | 2 | 2 | 2 | 2 | 2 | 0.5 (S) | 0.12 (S) | 1 (S) |
| *S. aureus* ATCC 29740 (Newbould) | 4 | 2 | 2 | 1 | 2 | 4 | 0.25 (S) | nd | 0.5 (S) |
| *S. aureus* ATCC BAA-41, MRSA | 4 | 2 | 2 | 2 | 2 | 1 | >32 (R) | >16 (R) | 1 (S) |
| *S. aureus* ATCC 700699, MRSA, Mu50 | 4 | 2 | 2 | 2 | 2 | 2 | >32 (R) | >16 (R) | 8 (I) |
| *Staphylococcus epidermidis* ATCC 12228 | 4 | 2 | 2 | 2 | 2 | 2 | 0.25 (S) | nd | 1 (S) |
| *S. saprophyticus* ATCC 15305 | 2 | 2 | 2 | 1 | 2 | 1 | 0.25 (S) | nd | 1 (S) |
| *S. haemolyticus* CDC HIP-5979, MRCoNS | 2 | 2 | 2 | 2 | 2 | 1 | >16 (R) | nd | 8 (I) |
| *S. hominis* CDC M270 | 1 | 2 | 2 | 1 | 2 | 2 | 0.12 (S) | nd | 0.5 (S) |
| *Enterococcus faecalis* ATCC 29212 | 4 | 2 | 2 | 1 | 2 | 2 | 0.25 (S) | nd | 2 (S) |
| *E. faecium* ATCC 35667 | 2 | 2 | 1 | 2 | 1 | 1 | 1 (S) | nd | 2 (S) |
| *E. faecium*, VanA | 2 | 2 | 1 | 1 | 1 | 1 | 1 (S) | nd | >16 (R) |
| *Streptococcus pyogenes* ATCC 19615, Group A | 8 | 4 | 2 | 2 | 4 | - | 0.25 (S) | nd | nd |
| *S. agalactiae* 60440, Group B | 8 | 4 | 2 | 2 | 8 | 2 | 1 (S) | nd | nd |
| *Bacillus cereus* ATCC 11778 | 16 | 2 | 4 | 4 | 4 | 4 | nd | nd | 1 (S) |
| *B. subtilis* ATCC 6633 | 16 | 2 | 2 | 2 | 4 | 4 | nd | nd | 0.25 (S) |
| *B. subtilis* ATCC 23857 | 4 | 2 | 2 | 2 | 2 | 2 | nd | nd | 0.25 (S) |
| *Listeria monocytogenes* ATCC 13932 | 8 | 2 | 2 | 2 | 1 | 1 | 1 (S) | nd | 2 (S) |

Abbreviations: ATCC, American Type Culture Collection; CDC, Center for Disease Control and prevention strain collection; MRSA, Methicillin-resistant *S. aureus*; MRCoNS, methicillin-resistant coagulase-negative *Staphylococcus*; VanA, vancomycin-resistance element; Group A or B, Lancefield types; CIP, ciprofloxacin; MEM, meropenem; VAN, vancomycin; Breakpoints for resistance are from CLSI, S, susceptible; I, intermediate; R, resistant; nd, not done.

Next, we explored the spectrum of activity of the **TM**-compounds against Gram-negative bacteria, which are notoriously less susceptible to antibiotic action. We tested the activity against a panel of *E. coli* strains that cause difficult-to-treat infections, namely, strains that possess a carbapenemase (e.g., NDM-1), various extended-spectrum β-lactamases, strains that are colistin resistant, ciprofloxacin resistant, or tetracycline resistant (**Table 3**). Most of the strains tested were multi-resistant, and their susceptibility profiles toward traditional drugs are also reported. **TO** analogs containing aromatic substituent at the C-3 appendage outperformed **TM-02** in terms of activity, with most of their MIC values ranging from 8 to 16 µg/ml against these antibiotic-resistant isolates of *E. coli*. Changing the halogen substituent of **TM-184** did not result in a substantial improvement in growth inhibitory activity (MIC of **TM-184** *vs* that of **TM-218** or **TM-220**), as observed for Gram-positive bacteria (**Table 2**).

**Table 3.** *In vitro* activity profile of TM-analogs against characterized antibiotic-resistant *Escherichia coli* isolates.

| *Escherichia coli* | Phenotype/<br>Genotype | Broth microdilution MIC (µg/ml) |  |  |  |  |  |  |  |  |  |  |  |  |
| --- | --- | --- | --- | --- | --- | --- | --- | --- | --- | --- | --- | --- | --- | --- |
|  |  | TM-02 | TM-184 | TM-218 | TM-220 | TM-247 | TM-303 | CAZ | MEM | CIP | CST | NIT | TET | TMP-S |
| ATCC 25922 | CLSI QC | 32 | 4 | 8 | 8 | 8 | 8 | 0.12 (S) | 0.06 (S) | 0.03 (S) | 1 (S) | 8 (S) | 1 (S) | 0.5/9.5 (S) |
| ATCC 35218 | TEM-1 | 32 | 16 | 8 | 16 | 16 | 8 | 0.25 (S) | 0.06 (S) | 0.25 (S) | 2 (S) | 0.25 (S) | 0.25 (S) | 0.12/2.3 (S) |
| ATCC BAA2471 | CRE; NDM-1 | 32 | 16 | 16 | 16 | 16 | 8 | >128 (R) | >32 (R) | >32 (R) | 1 (S) | 32 (S) | 128 (R) | >8/>152 (R) |
| 95882 CW-2011 | CRE, KPC | 32 | 16 | 16 | 16 | 16 | 8 | 128 (R) | 4 (R) | 32 (R) | 0.5 (S) | 8 (S) | 1 (S) | >8/>152 (R) |
| 70122 CW-2007 | ESBL; CTX-15M | 16 | 16 | 16 | 8 | 8 | 8 | 64 (R) | 0.06 (S) | >32 (R) | 1 (S) | 32 (S) | nd | >8/>152 (R) |
| ec302 | ESBL; TEM-10 | 32 | 16 | 16 | 16 | 16 | 32 | >128 (R) | 0.06 (S) | 0.06 (S) | 0.5 (S) | 16 (S) | 2 (S) | 0.5/9.5 (S) |
| ec305 | ESBL; SHV-1 | 4 | 2 | 2 | 4 | 4 | 2 | 1 (S) | 0.06 (S) | >32 (R) | 0.5 (S) | 8 (S) | >128 (R) | 1/19 (S) |
| ec308 | ESBL; SHV-5 | 8 | 4 | 4 | 2 | 4 | 2 | 128 (R) | 0.06 (S) | 0.25 (S) | 0.5 (S) | 8 (S) | 128 (R) | 0.12/2.3 (S) |
| ec309 | ESBL; MIR-1 | 32 | 16 | 16 | 16 | 16 | 8 | 64 (R) | 0.06 (S) | 0.06 (S) | 1 (S) | 16 (S) | 2 (S) | 0.5/9.5 (S) |
| ec310 | ESBL; ACT-1 | 32 | 16 | 16 | 16 | 16 | 32 | 64 (R) | 0.06 (S) | 0.06 (S) | 1 (S) | 8 (S) | 2 (S) | 0.25/4.7 (S) |
| ec125 | NIT-R | 32 | 32 | 16 | 16 | 32 | 16 | 1 (S) | 0.06 (S) | >32 (R) | 0.25 (S) | 128 (R) | 128 (R) | >8/>152 (R) |
| ec261 | TET-R; *tetM* | 32 | 4 | 4 | 4 | 4 | 2 | 0.25 (S) | 0.06 (S) | 2 (I) | 1 (S) | 4 (S) | 32 (R) | 1/19 (S) |
| 94474 CW-2010 | CST-R; *mcr-1* | >64 | 16 | 8 | 8 | 16 | 8 | 0.25 (S) | 0.06 (S) | 32 (R) | 32 (R) | 64 (I) | 128 (R) | >8/>152 (R) |
| 92886 CW-2010 | CST-R | 32 | 8 | 8 | 8 | 8 | 4 | 0.25 (S) | 0.06 (S) | 0.06 (S) | 8 (R) | 16 (S) | 1 (S) | 0.5/9.5 (S) |
| 93341 CW-2010 | CST-R | 32 | 4 | 8 | 8 | 8 | 2 | 0.25 (S) | 0.06 (S) | 0.25 (S) | 4 (R) | 64 (I) | 1 (S) | 0.5/9.5 (S) |
| 117167 CW-2015 | CST-R | 32 | 8 | 8 | 8 | 8 | 4 | 0.25 (S) | 0.06 (S) | 0.06 (S) | 4 (R) | 16 (S) | 0.5 (S) | 0.5/9.5 (S) |

Abbreviations: ATCC, American Type Culture Collection; CW-, CANWARD strain collection; CRE, carbapenem-resistant *Enterobacteriaceae*; ESBL, extended spectrum β-lactamase producer; TEM-1, TEM-10, NDM-1, KPC, CTX-15-M, MIR-1, ACT-1, SHV-1, and SHV-5 are β-lactamase names; TET, tetracycline; CAZ, ceftazidime; MEM, meropenem; CIP, ciprofloxacin; CST, colistin; NIT, nitrofurantoin; TMP-S, trimethoprim-sulfamethoxazole; Breakpoints for resistance are from CLSI, S, susceptible; I, intermediate; R, resistant; nd, not done.

Surprisingly, **TM-184** (iodine) displayed greater bactericidal activity (≥3 log10 decrease in CFU/ml within 24 h) than **TM-218** (bromine) or **TM-220** (chlorine) against the carbapenem-resistant *E. coli* strain (BAA 2471, NDM-1). At 1× or 2×MIC, **TM-184** showed rapid and more extensive killing than the two other derivatives (**Fig. 1**). This was also generally true for the bactericidal activity of **TM-184** against the extremely resistant (XDR) *Acinetobacter baumannii* strain BAA-1800, also a carbapenem-resistant strain, although in this case the difference between **TM-184** and **TM-218** was minimal against that bacterial species (**Fig. 2**). As seen for *E. coli*, these **TO** derivatives had the same MIC (16 µg/ml) against the two *A. baumannii* strains tested (**Table 4**). Noteworthy, we could not isolate resistant colony from kill kinetics studies (**Fig. 1 and Fig. 2**), even though some regrowth was observed in cultures containing the lowest drug concentration. We thus presumed the regrowth was caused by compound precipitation or instability.

**Fig. 1.**
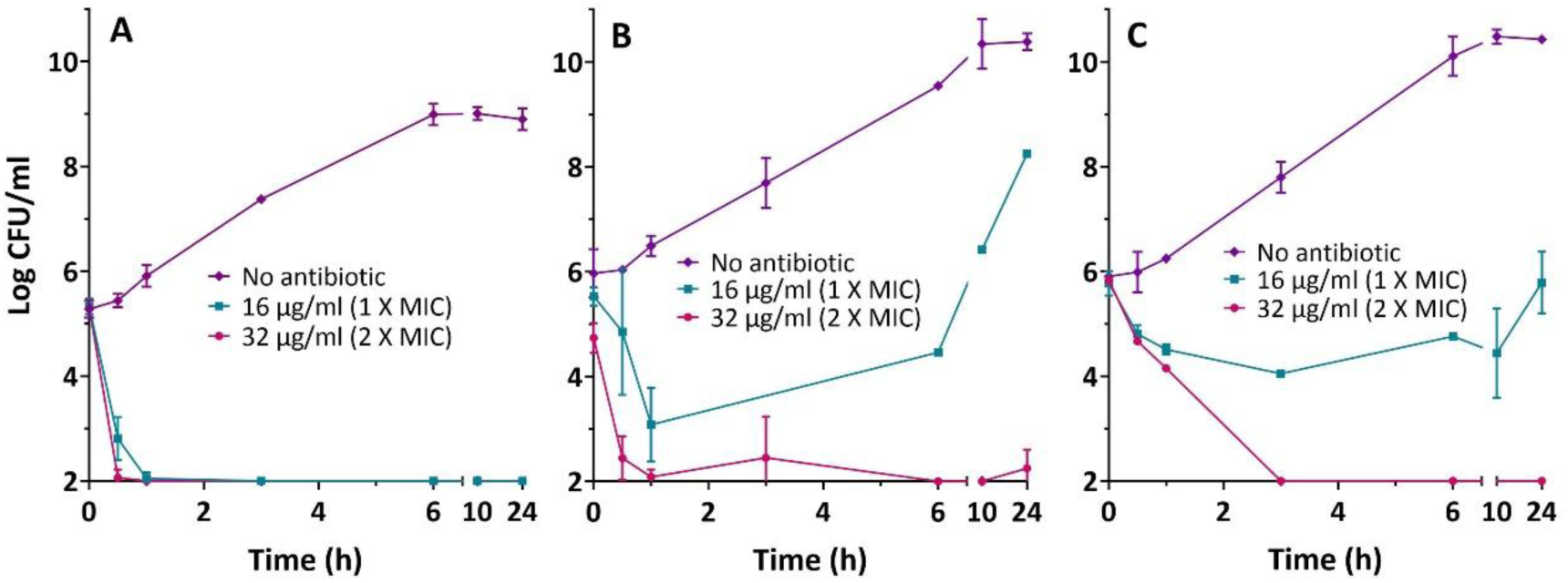
Bactericidal effects of TM-184 (A), TM-218 (B), and TM-220 (C) on *Escherichia coli* ATCC BAA 2471 (CRE, NDM-1). Kill kinetics were assessed as described in the Methods and Materials section. The log CFU/ml at each time point represents the mean of three independent experiments. The detection limit was 100 CFU/ml.

**Fig. 2.**
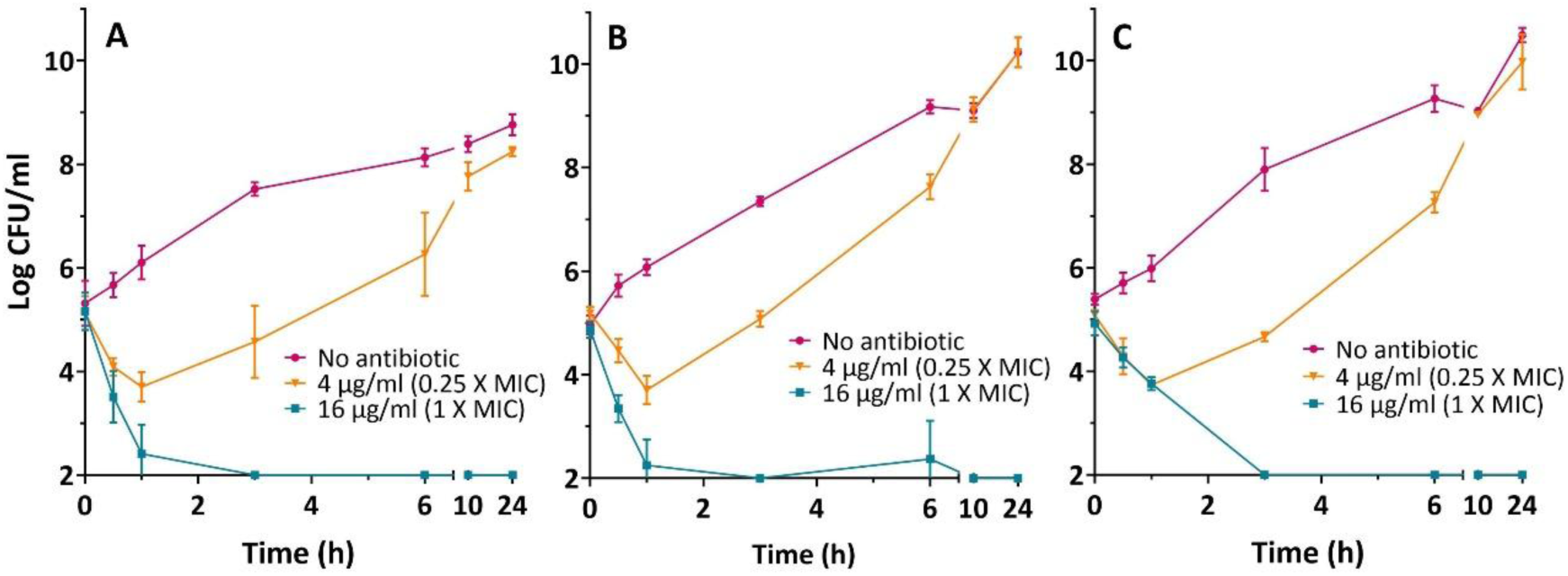
Bactericidal effects of TM-184 (A), TM-218 (B), and TM-220 (C) on *Acinetobacter baumannii* ATCC BAA 1800 (XDR). Kill kinetics were assessed as described in the Methods and Materials section. The log CFU/ml at each time point represents the mean of three independent experiments. The detection limit was 100 CFU/ml.

**Table 4.** *In vitro* activity profile of TM-compounds against antibiotic-resistant *Acinetobacter baumannii*.

|  | Broth microdilution MIC (µg/ml) |  |  |  |  |  |  |  |  |  |  |
| --- | --- | --- | --- | --- | --- | --- | --- | --- | --- | --- | --- |
|  | TM-02 | TM-184 | TM-218 | TM-220 | TM-247 | TM-303 | CAZ | CIP | GEN | MEM | TET |
| *A. baumannii* |  |  |  |  |  |  |  |  |  |  |  |
| ATCC 19606 | >64 | 16 | 8 | 8 | 16 | 8 | 8 (S) | 0.5 (S) | 32 (R) | 2 (S) | 2 (S) |
| ATCC BAA-1800, XDR | >64 | 16 | 16 | 16 | 16 | 16 | 128 (R) | >32 (R) | >128 (R) | >32 (R) | 128 (R) |

Abbreviations: ATCC, American Type Culture Collection; XDR, extreme drug resistance; CAZ, ceftazidime; CIP, ciprofloxacin; GEN, gentamicin; MEM, meropenem; TET, tetracycline; Breakpoints for resistance are from CLSI, S, susceptible; I, intermediate; R, resistant.

These results prompted us to test the activity of **TM-184** against an extended panel of Gram-negative species (**Table 5**). **TM-184** showed MIC ranging from 8 to 16 µg/ml against *Haemophilus influenzae*, *Shigella* and *Citrobacter* strains. However, most other *Enterobacterales* such as *Proteus*, *Serratia*, *Salmonella*, *Enterobacter* and *Klebsiella* were not susceptible to the drug (MIC ≥32 µg/ml). This was also true for multi-resistant *Pseudomonas aeruginosa* strains (MIC ≥64 µg/ml), even though the CLSI antibiotic-quality control strain ATCC 27853 was relatively susceptible to **TM-184** (MIC 8 µg/ml).

**Table 5.** *In vitro* activity profile of TM-184 against a variety of Gram-negative bacterial species.

| Bacterial strains | Broth microdilution MIC (µg/ ml) |  |  |
| --- | --- | --- | --- |
|  | TM-184 | CIP | IMI |
| *Haemophilus influenzae* ATCC 49766 | 8 | ≤0.03 (S) | nd |
| *Haemophilus influenzae* ATCC 49247 | 16 | ≤0.03 (S) | nd |
| *Shigella flexneri* ATCC 12025 | 8 | ≤0.03 (S) | 0.5 (S) |
| *Citrobacter freundii* ATCC 8090 | 8 | ≤0.03 (S) | 1 (S) |
| *Enterobacter cloacae* ATCC 35030 | 32 | ≤0.03 (S) | 0.06 (S) |
| *Enterobacter cloacae* ATCC BAA 2468, CRE (NDM-1) | 64 | nd | >16 (R) |
| *Klebsiella oxytoca* ATCC 43165 | 16 | ≤0.03 (S) | 1 (S) |
| *Klebsiella oxytoca* ATCC 13182 | 32 | ≤0.03 (S) | 1 (S) |
| *Klebsiella aerogenes* ATCC 35029 | 64 | ≤0.03 (S) | 1 (S) |
| *Klebsiella pneumoniae* ATCC 13883 | 32 | ≤0.03 (S) | 1 (S) |
| *Klebsiella pneumoniae* ATCC BAA 2473, CRE (NDM-1) | >64 | >32 (R) | >16 (R) |
| *Klebsiella pneumoniae* ATCC BAA 1705, ESBL | >64 | >32 (R) | >16 (R) |
| *Proteus mirabilis* ATCC 25933 | >64 | ≤0.03 (S) | 0.12 (S) |
| *Serratia marcescens* ATCC 8100 | >64 | 0.06 (S) | 0.5 (S) |
| *Salmonella typhimurium* 8375, FFC-R | >64 | ≤0.03 (S) | 1 (S) |
| *Salmonella typhimurium* 15789, FFC-R | >64 | ≤0.03 (S) | 1 (S) |
| *Salmonella enterica* ATCC 14028 | >64 | ≤0.03 (S) | 1 (S) |
| *Salmonella enteritidis* ATCC 4931 | >64 | ≤0.03 (S) | 1 (S) |
| *Pseudomonas aeruginosa* ATCC 27853 | 8 | 0.25 (S) | 1 (S) |
| *Pseudomonas aeruginosa* ATCC BAA-2108, XDR | >64 | 1 (S) | 8 (R) |
| *Pseudomonas aeruginosa* ATCC BAA-2114, XDR | 64 | 2 (I) | 1 (S) |
| *Pseudomonas aeruginosa* 91196, GES | >64 | 128 (R) | >16 (R) |
| *Pseudomonas aeruginosa* 103843, VIM | >64 | >128 (R) | >16 (R) |
| *Pseudomonas aeruginosa* 96918, IMP | >64 | >128 (R) | >16 (R) |

Abbreviations: ATCC, American Type Culture Collection; XDR, extreme drug resistance; CRE, carbapenem-resistant *Enterobacteriaceae*; ESBL, extended spectrum β-lactamase producer; FFC-R, florfenicol-resistant; NDM-1, GES, VIM, and IMP are β-lactamase names; CIP, ciprofloxacin; IMI, imipenem. Breakpoints for resistance are from CLSI, S, susceptible; I, intermediate; R, resistant.

2.4. *In vivo* activity

Compound **TM-184** with a MIC 4-8 µg/ml against *E. coli* ATCC 25922 was next evaluated in the neutropenic mouse, thigh infection model, using cefotaxime as a comparator drug (MIC 0.12 µg/ml). Mice were infected with *E. coli* ATCC 25922, and compounds solubilized in 45% cyclodextrin were administered by 2 intraperitoneal injections of 15 mg/kg of body weight at 2 and 6 hours after infection (total dose 30 mg/kg). **TM-184** showed *in vivo* activity, causing a significant reduction of the bacterial load (**Fig. 3**). The treatment control drug cefotaxime was also used at the same dose of 30 mg/kg (a relative dose of 250-times its MIC, *i.e*., a dose of 30/MIC of 0.12) and was accordingly more active than **TM-184** used at 7.5-times its MIC (*i.e*., a dose of 30/MIC of 4).

**Fig. 3.**
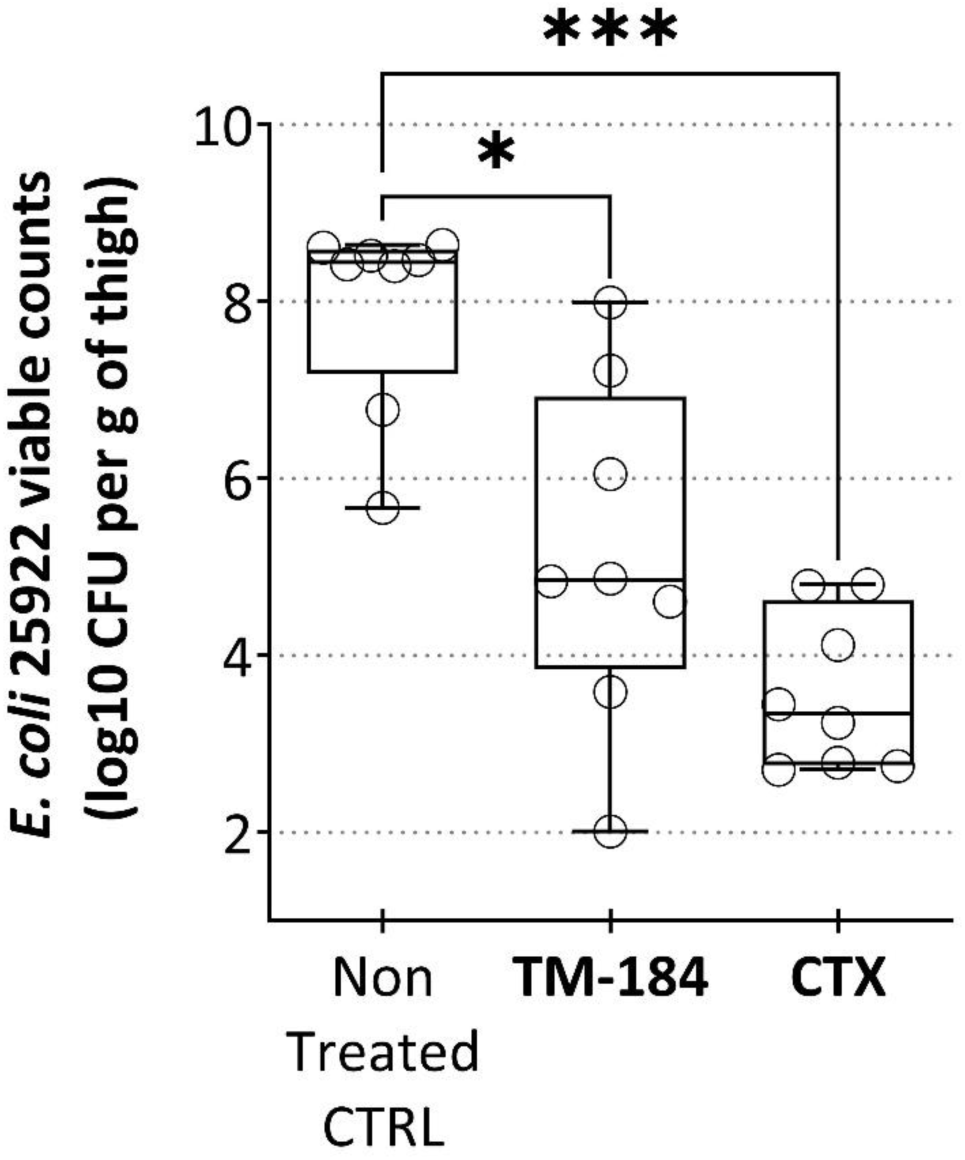
Activity of **TM-184** in a neutropenic mouse thigh infection model. Mice were infected with *E. coli* ATCC 25922, and **TM-184**, dissolved in 45% cyclodextrin, was administered intraperitoneally (IP) at 15 mg/kg, 2 and 6 h post-infection. Cefotaxime (CTX) was used as a treatment control (also IP, same dose, same diluent). Horizontal bars within the boxes indicate the median value for each group, and whiskers show the minimum and maximum values. Results were compared with the placebo control group, which received only the diluent (non-treated CTRL). Both compounds significantly reduced bacterial loads *in vivo*. Statistical significance was determined using the Kruskal–Wallis test followed by Dunn’s multiple-comparison test (* p < 0.05, *** p < 0.001).

## 3. Discussion

In a recent structure-activity relationship (SAR) study, we investigated C3-derivatives of tomatidine (**TO**) [14]. This extensive research was facilitated by the development of a novel, gram-scale synthesis of **TO** [19,20], leading to the discovery of the analog **TM-02**, which showed modest activity (MIC of 32 µg/ml) against the Gram-negative bacterium *Escherichia coli*. To further enhance antibiotic activity against Gram-negative bacteria, the SAR of **TO** and **TM-02** steroid analogs generated nearly 100 molecules with various modifications, including alkyl, cyclic, and (hetero)aromatic substituents. Aromatic substituents generally exhibited better antibacterial profiles, with **TM-184** emerging as a leading compound due to its C3-modification, which allowed it to bypass the outer membrane barrier and achieve a 4- to 8-fold improvement in activity against *E. coli* ATCC 25922 [14]. The antibacterial activity of **TM-184** against *E. coli* ATCC 25922 (MIC 4 µg/ml) was significantly better than that of **TO** (MIC >128 µg/ml) and **TM-02** (MIC 32 µg/ml). This study also demonstrated that **TM-184** has a high affinity for *E. coli* ATP synthase (Ki 0.8-1.3 µM), similar to **TM-02** [16], and that its increased activity compared to **TM-02** was mainly due to improved whole-cell permeability (see permeability ratio, **Table 1**). Intrinsic resistance due to efflux was low for **TM-184** (see efflux ratio, **Table 1**), and deletion of the *E. coli* AcrAB pump had only a minor effect (drop of one MIC dilution). However, this small efflux pump effect may perhaps explain the slight reduction in **TM-184**’s activity against many clinical *E. coli* MDR strains (**Table 3**), which often exhibit resistance to multiple antibiotic classes due to efflux pump overexpression [21]. It remains to be determined whether both the permeability barrier and efflux contribute to the poor activity of **TM-184** against *Klebsiella spp*. and *Pseudomonas aeruginosa* (**Table 5**). Conversely, our series of C3-derivatives of **TO** showed consistent activity against various Gram-positive bacteria, including MRSA, where the ATP synthase target is readily accessible at the cell surface (**Table 2**).

In the SAR study of C3-derivatives of **TO**, Delbrouck *et al*. [14] demonstrated that substituting a phenyl ring at the C3 position with halides in the para position (*e.g*., **TM-184**, **TM-218**, **TM-220**) enhanced activity against *E. coli* strains by 4 to 8-fold, except for fluorine substitution, which had no effect. While these compounds were equally effective in terms of MIC (**Table 3**), our findings indicate that the iodine derivative (**TM-184**) exhibited significantly greater bactericidal activity against both *E. coli* and *Acinetobacter baumannii*. Previously, we showed that the bactericidal activity of **TO** was linked to increased production of reactive oxygen species following ATP synthase inhibition [10]. The rapid killing of the NDM-1 producing *E. coli* strain BAA-2471 and the carbapenem-, fluoroquinolone-, aminoglycoside-, and tetracycline-resistant *Acinetobacter* strain BAA-1800 by **TM-184** at the MIC was particularly noteworthy (**Fig. 1** and **Fig. 2**), as both bacteria are high-priority targets for new treatments [2]. *A. baumannii*, in particular, is known for causing extremely difficult-to-treat infections in hospital intensive care units, including pneumonia, wound, bloodstream, and urinary tract infections [3].

Fortunately, our inhibitors demonstrated significantly lower inhibitory activity against mitochondrial ATP synthase compared to oligomycin, a known ATP synthase inhibitor with mammalian toxicity [18]. Specifically, our compounds exhibited at least 800-fold lower inhibitory activity than oligomycin (**Table 1**). To follow this inquiry, it was legitimate to ask if a difference in mitochondrial ATP synthase inhibition compared to that provoked by oligomycin (or other known ATP synthase inhibitors like venturicidin) translates to different levels of cytotoxicity. Preliminary assessment of **TM-184** cytotoxicity for HepG2 cells (72-h exposure time) showed that **TM-184** was indeed much less cytotoxic than oligomycin and venturicidin (**Fig. S4**). Based on these findings, we proceeded to evaluate **TM-184** in mice to assess its *in vivo* activity. **TM-184** was able to significantly reduce the level of infection by *E. coli* in this model (**Fig. 3**). However, the variability in data points observed for **TM-184** in **Fig. 3** may be due to the limited aqueous solubility of this compound intrinsic to the steroid scaffold, suggesting a need for formulations that enhance bioavailability. Orienting a new SAR study to further improve both the cytotoxicity and pharmacological parameters of **TM-184** seems to be the next logical step in this research program.

There is strong evidence that C3-derivatives of **TO** target bacterial ATP synthase, with their antibacterial action closely linked to the inhibition of this enzyme [14, 16]. This is supported by a clear correlation between the whole-cell activity of C3-derivatives and their affinity (IC_50_) for the ATP synthase of *E. coli*, as well as *in silico* docking studies [14]. However, the partial retention of activity against an *S. aureus* SCV with a point mutation in *atpE* (**Table 1**) suggests that a secondary mechanism of action may contribute to the residual whole-cell activity observed in such mutants. Further investigation is needed to understand the significance of this observation in whole-cell activity.

## 4. Conclusion

Our previous work on expanding the activity spectrum of tomatidine (**TO**) through structure-activity relationship (SAR) studies of C3-modifications has successfully led to the identification of 4 interesting analogs of **TM-02** [14]. In this study, we further explored the antibacterial activity profile of **TM-184, TM-218, TM-220, TM-227**, and **TM-303**, highlighting **TM-184** as the standout candidate with potent and broad-spectrum bactericidal effects and *in vivo* efficacy against carbapenem-resistant Gram-negative bacteria, including NDM-1-producing *E. coli* and *Acinetobacter*. These findings demonstrate that broad-spectrum ATP-synthase inhibitors can be advanced as a new antibiotic class. By harnessing tomatidine and other steroid scaffolds, they directly address the WHO’s call for agents active against priority Gram-negative and Gram-positive pathogens, even though their complex architectures present notable synthetic and pharmacokinetic challenges.

## 5. Experimental

### 5.1. General Information

Compounds of this study are reported in Delbrouck *et al*. [14]. Analogs **TM-02**, **TM-184**, **TM-218**, **TM-220**, **TM-247**, and **TM-303** correspond to molecules #2, #41, #40, #39, #34, and #42, from the aforementioned article wherein synthetic protocols and structural characterizations (^1^H, ^13^C NMR, and High-resolution electrospray mass spectroscopy HRMS) are described. The known ATP synthase inhibitors 4-chloro-7-nitrobenzofurazan (NBD-Cl) and oligomycin were both from Sigma-Aldrich (Oakville, ON, Canada).

### 5.2. Bacterial strains and growth conditions

Bacterial strains labeled ATCC or BAA originated from the American Type Culture Collection (ATCC, Rockville, MD), while those labeled CDC came from the Centers for Disease Control and Prevention (Atlanta, GA). Several clinical strains were also sourced from the Canadian Ward Surveillance Study (CANWARD) [22], with additional strains from Malouin’s laboratory bacterial culture collection at Université de Sherbrooke, QC, Canada. SCV strains of *Staphylococcus aureus* NewbouldΔ*hemB* and Δ*hemB atpE* (SaR5-1), which possess a point mutation in AtpE (G18C) conferring tomatidine resistance, were previously documented [9]. Additionally, the parental *Escherichia coli* strain MC4100 (ATCC 35695), its membrane-compromised mutant strain MC4100 *lptD4213* (formerly known as imp4213 with an altered *lptD* gene) [23], and its efflux pump-deficient mutant MC4100 *acrAB* [24], were utilized to examine the effects of the outer membrane barrier and compound efflux on antibiotic susceptibility, as previously reported [14]. Tryptic soy broth (TSB; BD Biosciences, Mississauga, ON, Canada) or tryptic soy agar (TSA; BD Biosciences) was employed to maintain bacterial cultures, while other specified media were used for antibiotic susceptibility testing.

### 5.3. Antibiotic susceptibility testing

The minimum inhibitory concentrations (MIC) of antibiotics were assessed using a broth microdilution method, following the Clinical Laboratory and Standards Institute (CLSI) guidelines for antimicrobial susceptibility testing [25], as previously detailed [14]. Antibacterial agents were prepared in cation-adjusted Mueller-Hinton broth (CAMHB, BD Biosciences) and subjected to two-fold serial dilutions in 96-well plates. Each well received an equal volume of a bacterial suspension (∼10^5^-10^6^ CFU/ml) and was incubated for 18–24 h at 35°C for staphylococci or 37°C for other bacteria. Sterility and bacterial growth controls were included in all assays. *Staphylococcus aureus* ATCC 29213 and *Escherichia coli* ATCC 25922 served as quality control strains for all antibiotic susceptibility tests. Reference drugs or test performance controls included ceftazidime, ciprofloxacin, colistin, meropenem, imipenem, nitrofurantoin, tetracycline, gentamicin, trimethoprim-sulfamethoxazole, and/or vancomycin (all from Sigma-Aldrich). The MIC was defined as the lowest concentration of the drug that showed no visible growth. MIC determinations were conducted at least three times, with the reported values representing the most frequent result. Breakpoints for resistance (R), intermediate resistance (I), and susceptibility (S) for reference drugs were based on CLSI guidelines [25].

### 5.4. Kill kinetics

Bacteria were inoculated at approximately 10^5^-10^6^ CFU/ml in CAMHB, with or without antibiotics at the specified concentrations. During growth at 37°C (225 RPM), samples were taken at various time points, serially diluted, and plated on TSA (18-24 h at 37°C) to determine CFU (viable bacterial counts).

### 5.5. Bacterial ATP synthase assay

#### 5.5.1. Membrane vesicles (MV) as the source of bacterial ATP synthase

The method for preparing MV was adapted from [9] and previously described in [14]. *E. coli* ATCC 25922 and *S. aureus* ATCC 29213 served as the sources of MV. The MV were suspended in a solution of 50 mM MOPS-10 mM MgCl2 MOPS with 10% glycerol, aliquoted, and stored at −80°C. The protein content of MV was determined using the Micro BCA protein assay kit (Thermo Fisher, Rockford, IL).

#### 5.5.2. ATP production by MV and IC_50_ determination

As previously described [14], MV (30 µg proteins/ml) were incubated with various concentrations of inhibitors for 30 min. ATP synthesis was initiated by adding 2.5 mM NADH and 1 mM ADP in a 10 mM potassium phosphate buffer (final concentrations). The reaction was allowed to proceed for 60 min before being stopped with 2 mM ethylenediaminetetraacetic acid and 1% trichloroacetic acid. A portion of the reaction mixture was transferred to white 96-well plates, and 50 μL of a luciferin/luciferase reagent (ATPlite one-step Luminescence Assay System kit; PerkinElmer, Woodbridge, ON, Canada) was added. This reagent oxidizes luciferin to oxyluciferin, emitting light proportional to the ATP amount present. The signal, indicating the ATP produced by the ATP synthase of the MV, was measured using a luminometer (LumiStar OPTIMA microplate reader; BMG Labtech, Cary, NC). The inhibitor concentration that prevented 50% of ATP production by the ATP synthase (IC_50_) was determined using GraphPrism software (v 9.2.0), employing the log(inhibitor) *vs*. response-variable slope (four parameters) tool, and expressed in µM. Each IC_50_ determination was the mean (and standard deviation) of two or mostly three independent experiments using at least 7 to 12 different inhibitor concentrations, resulting in a response-variable slope of r^2^ ≥0.9.

#### 5.5.3. Determination of the inhibitor constant Ki

Enzyme kinetics assays were conducted to obtain precise information about the Ki, which is the concentration of inhibitor needed to achieve half-maximal inhibition. ATP production by the MV was measured as described in section (ii), with a few modifications. The pre-exposure time for the inhibitors during the kinetics tests ranged from 0 to 10 min. Various reaction times were tested to determine the enzyme’s rate, and different substrate concentrations were used to assess the enzyme’s affinity for its substrate. These tests allowed for the determination of the maximum rate of ATP production (Vmax) and the affinity constant for the substrate (Km). Specifically, inhibition kinetics data were analyzed to determine the inhibition constants (Ki) using the method of Burlingham and Widlanski [26], as previously described [16]. In brief, the inverse of the luminescence production rate (s/RLU) was plotted against the inhibitor concentration. The y-intercept value was then divided by the slope. To obtain the Ki, this value was corrected with the Km, which was calculated by measuring ATP production with varying ADP concentrations (0.05-1 mM) over a 25-min period under similar conditions as described above.

### 5.6. Mitochondrial ATP synthase assay

#### 5.6.1. Preparation of mitochondria

Hep-2 cells (ATCC CCL-23) were used as the source of mitochondria, which were obtained exactly as described before [14]. The protein content of the mitochondria samples was assayed by the Pierce Rapid Gold BCA Protein Assay Kit (Thermo Fisher Scientific, Rockford, IL).

#### 5.6.2. Mitochondrial ATP production and inhibition (IC_50_ determination)

As previously described [14], mitochondria were diluted in an assay buffer (Tris-HCl, KH_2_PO_4_, sorbitol, MgSO_4_, bovine serum albumin) to achieve a final protein concentration of 150 µg/ml. Various concentrations of inhibitors were added to the mitochondrial suspension and incubated for 30 min. To initiate ATP synthesis, 5 mM succinate and 67 µM ADP (final concentrations) were added. The reaction was halted after 60 min by adding 1.75 µL of 60% perchloric acid and placing the reaction mixture on ice for 10 min. This was followed by centrifugation at 15,300 × *g* for 10 min. The supernatant was transferred to another tube containing 11.5 µL of 1 M KOH to neutralize the acid. The mixture was incubated on ice for 3 min and centrifuged again. A small volume of the supernatant was transferred to a white 96-well plate. The plate was read on a luminometer after adding 50 μL of the luciferin/luciferase reagent from the ATPlite one-step Luminescence Assay System kit, as described for bacterial MV. The IC_50_, or the concentration of inhibitor that prevents 50% of ATP production by mitochondrial ATP synthase, was determined using GraphPrism software, as previously described.

### 5.7 Preliminary cytotoxicity assay

Cell viability was measured using the Cell Proliferation Kit I from Roche (Sigma-Aldrich, Oakville, ON, Canada). In this assay, the conversion of 3-(4,5-dimethylthiazol-2-yl)-2,5-bromide diphenyltetrazolium (MTT) to formazan crystals is correlated with cellular activity. The day prior to the cytotoxicity assay, HepG2 cells (ATCC HB-8065) were seeded in 96-well plates (2 × 104 cells/well) and grown for 24 h. Then, cells were exposed to test compounds at various concentrations in Dulbecco modified Eagle medium (DMEM; Wisent, Saint-Jean-Baptiste, QC, Canada) containing 1% fetal bovine serum (FBS; Wisent), GlutaMAX (Gibco, Thermo Scientific, St-Laurent, QC, Canada) and 4.5 g of glucose per liter. Cells were incubated for 24 h or 72 h at 37°C in 5% CO_2_ atmosphere with 95% humidity. At the end of the drug exposure period, 500 mg/L of MTT was added to each of the wells, and the plates were incubated for an additional 4 h at 37°C in 5% CO_2_ atmosphere with 95% humidity. A solubilizing solution (10 % sodium dodecyl sulfate in 0.01 M hydrochloric acid) was added to each well, and the cellular reduction of MTT (yellow tetrazolium salt) and production of formazan (purple crystals) was measured after 24 h with a scanning multiwall spectrophotometer (550 nm/ 690 nm). A low amount of formazan indicated a loss of cell viability. Inhibitory concentrations 50% (IC50) were derived from the dose-response curve for each compound and were calculated with the GraphPrism software, employing the log(inhibitor) vs. response-variable slope (four parameters) tool.

### 5.8. Neutropenic mouse thigh infection model

Female CD-1 mice (Charles River, Sherbrooke, QC, Canada), weighing 22-25 g, were immunosuppressed with two cyclophosphamide injections: 150 mg/kg four days prior and 100 mg/kg one day before the infection challenge. On the day of infection, mice were anesthetized with a ketamine (87 mg/kg) and xylazine (13 mg/kg) mixture. Each thigh (quadriceps) was then inoculated with approximately 5 × 10^5^ *E. coli* ATCC 25922 in a 50-µL volume. To assess activity, compounds were administered intraperitoneally at 15 mg/kg in a 200-µL volume, 2 and 6 h post-infection. The compounds were dissolved in 45% cyclodextrin. At 8 h post-infection, the mice were humanely euthanized, and the infected thigh tissues were collected, homogenized in PBS, and bacterial CFU per gram of tissue were determined by serial dilutions on TSA.

### 5.9. Statistical analysis

Statistical analyses and IC_50_ values were calculated using GraphPad Prism software (v.9.3.1). The specific statistical tests applied to the animal experimentation results were also conducted using GraphPad software and are detailed in the legend of the corresponding figure.

## Ethical approval

The animal studies adhered to the Canadian Council on Animal Care guidelines and received approval from the institutional ethics committee on animal experimentation at the Faculté des Sciences, Université de Sherbrooke (protocol code 2019-2252, approved on October 13, 2021).

## Declaration of generative AI and AI-assisted technologies in the writing process

During the preparation of this work the author(s) used Microsoft 365 Copilot in order to correct grammatical errors. No changes in results, interpretations, contents or cited references were made. After using this tool, the author(s) reviewed and edited the content as needed and take(s) full responsibility for the content of the publication.

