## Supplemental data for "*In vitro* and *in vivo* activity profiles of broad-spectrum bacterial ATP synthase inhibitors"

### **Corresponding authors:**

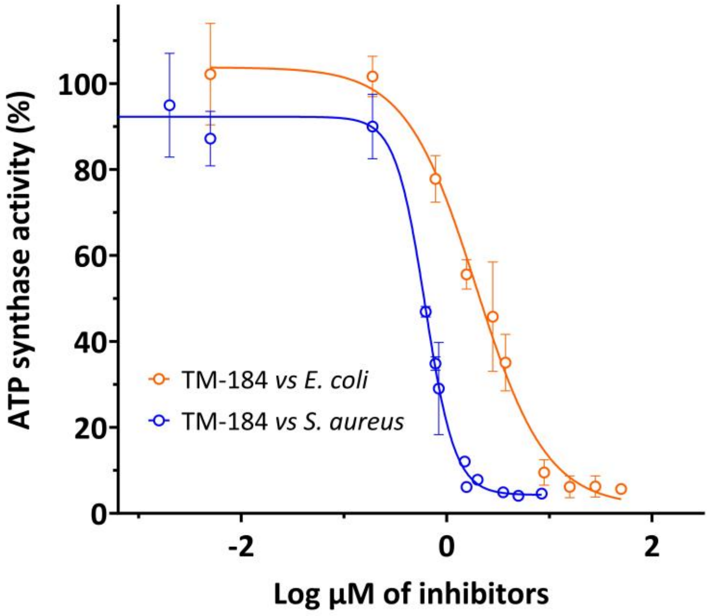


**Figure S1.** Illustration of experimental data for determining the $IC_{50}$ of TM-184 using membrane vesicles from *Staphylococcus aureus* (blue) and *Escherichia coli* (orange) as sources of bacterial ATP synthases. The luminescent signal, indicating the amount of ATP produced by the membrane vesicles, was measured using a luciferin/luciferase assay system, as detailed in the Methods and Materials section. The $IC_{50}$ was calculated with GraphPrism software, employing the log(inhibitor) *vs*. response-variable slope (four parameters) tool, and expressed in µM. The $r^2$ values for the *S. aureus* and *E. coli* curves were 0.9816 and 0.9660, respectively. The $IC_{50}$ values are reported in the results section (Table 1).

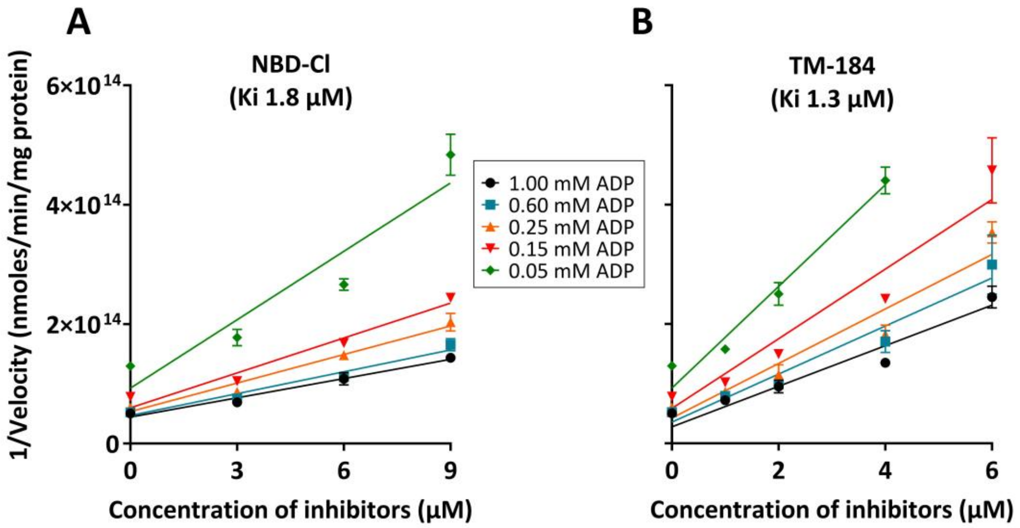


**Figure S2.** Here is an example of experimental data for determining the Ki of compounds NBD-Cl (A) and TM-184 (B) using membrane vesicles from *Escherichia coli* as the source of bacterial ATP synthase. The luminescent signal, indicating the amount of ATP produced by the membrane vesicles, was measured using a luciferin/luciferase assay system, as detailed in the Methods and Materials section. Various substrate (ADP) concentrations were tested to assess the enzyme's affinity for its substrate. The Ki values were determined as described in the Methods and Materials section and are reported in Table 1 (see text, results section).

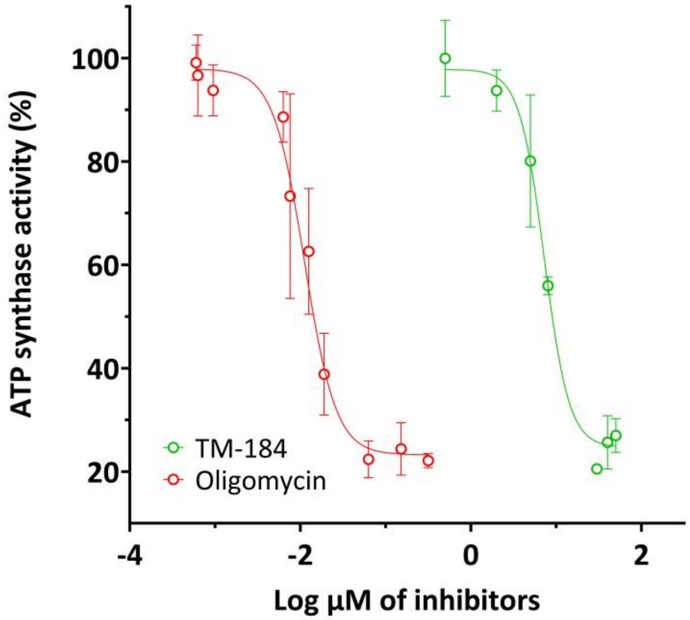


**Figure S3.** Here is an example of experimental data for determining the $IC_{50}$ of TM-184 and oligomycin using mitochondria isolated from HEp-2 cells as the source of bacterial ATP synthase. The luminescent signal, indicating the amount of ATP produced by the membrane vesicles, was measured using a luciferin/luciferase assay system, as detailed in the Methods and Materials section. The $IC_{50}$ was calculated with GraphPrism software, employing the log(inhibitor) *vs*. response-variable slope (four parameters) tool, and expressed in µM. The $r^2$ values for the TM-184 and oligomycin curves were 0.9635 and 0.9156, respectively. The relative $IC_{50}$ of TM-184 was 845-fold higher than that of oligomycin (Table 1).

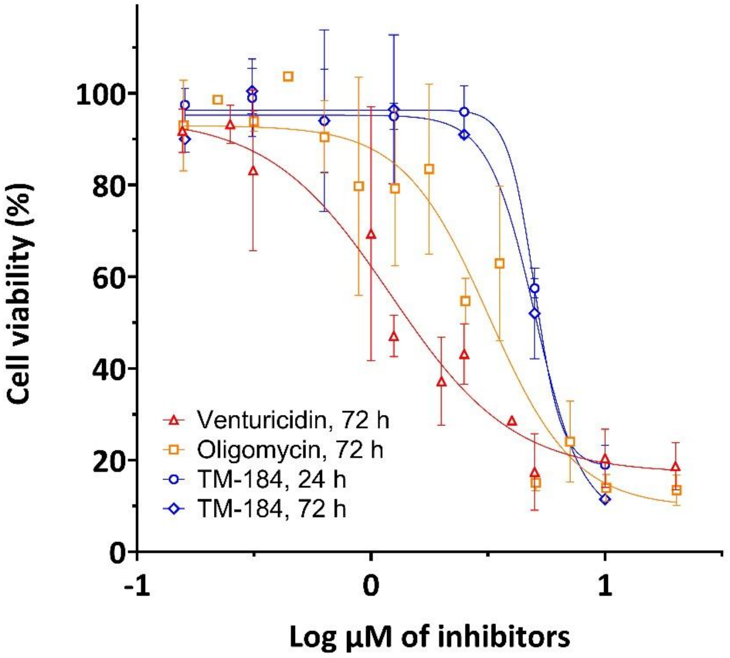


**Figure S4.** As shown in previous Figure S3, the relative $IC_{50}$ of TM-184 for the mitochondrial ATP synthase was 845-fold higher than that of oligomycin (Table 1). To follow this inquiry, it was legitimate to ask if a difference in mitochondrial ATP synthase inhibition compared to that provoked by oligomycin (or other known ATP synthase inhibitors like venturicidin) translates to different levels of cytotoxicity. To answer this question, we exposed HepG2 cells to increasing concentration of those inhibitors for 24 h or 72 h and measured cell viability using a MTT assay (Sigma-Aldrich, Oakville, ON, Canada). The $IC_{50}$ was calculated with GraphPrism software, employing the log(inhibitor) *vs*. response-variable slope (four parameters) tool, and was expressed in µM. The $r^2$ values for the TM-184 24 h and 72 h curves were 0.9732 and 0.9333, respectively, and were 0.8814 and 0.8962 for the oligomycin and venturicidin curves after 72 h. The relative $IC_{50}$ of TM-184 was 5.0 µM at 24 h or 72 h, and that of oligomycin and venturicidin at 72 h were 3.18 and 1.22 µM, respectively.
